# Acellular normothermic spleen perfusion resolves transcriptional and non-transcriptional mechanisms of steroid immunosuppression

**DOI:** 10.64898/2026.05.16.725632

**Authors:** Nicholas A. Callais, Shawn Welch, Randall R. Rainwater, John Schaller, Shuoqiu Deng, Ana Azevedo-Pouly, Marie Schluterman Burdine, Lyle Burdine

## Abstract

The limiting step in immune-active drug development is increasingly not candidate generation but testing whether a candidate therapy is effective in a system that preserves tissue architecture, vascular exposure, multicellular interaction, and repeated pharmacodynamic sampling without patient exposure. We developed an acellular normothermic machine-perfusion platform for intact porcine spleen designed as a translational immune-organ assay. Across independent acellular perfusions, the circuit maintained physiologic parameters, preserved red- and white-pulp histology, and yielded viable effluent cells suitable for serial flow cytometry and multiomics. High-dose methylprednisolone was used as a clinically familiar perturbation to determine whether the platform could resolve steroid immunosuppression at mechanistically distinct levels. Effluent RNA-seq identified canonical glucocorticoid-responsive transcriptional programs, including DUSP1, FKBP5, PER1, DDIT4, SGK1, KLF9, ANXA1, NF-κB feedback regulators, and JAK/STAT suppressor pathways. SOCS3 was a prominent early signal in the perfusion transcriptome and was validated orthogonally at the protein level in prednisone-treated, CD3/CD28-activated primary murine splenocytes, strengthening its role as a candidate pharmacodynamic marker. In parallel, data independent acquisition (DIA) proteomics of effluent cell pellets nominated a non-transcriptional protein-level response: a Sus scrofa LGALS13-annotated, CLC/Galectin-10-like galectin detected despite absence of the corresponding effluent-cell transcript. Because this porcine LGALS13-annotated protein group is treated here as an orthologous CLC/Galectin-10-like signal rather than as canonical human placental Galectin-13/PP13, we tested recombinant human Galectin-10 in vitro. Human Galectin-10 induced marked apoptosis of CD3/CD28-stimulated Jurkat cells, prioritizing this axis for future mechanistic testing without proving causality in the perfused spleen. These data establish acellular spleen perfusion as a serial multiomic platform for translational immunopharmacology and motivate deployment with otherwise-discarded human donor spleens.

**One sentence summary:** An acellular intact-spleen perfusion platform enables serial cellular, transcriptomic, proteomic, and functional pharmacodynamic sampling that identifies steroid-responsive transcriptional programs, validates SOCS3 protein induction, and nominates a CLC/Galectin-10-like non-transcriptional immunosuppressive axis for translation to discarded human donor spleens.

## Introduction

The translational bottleneck in drug development is changing. AI-enabled molecular design, high-throughput chemistry, structure-aware modeling, and multimodal omics now generate therapeutic hypotheses at a pace that conventional preclinical systems cannot absorb.^1–3^ This problem is especially acute for immune-active therapeutics, where efficacy and toxicity depend on vascular exposure, spatial compartmentalization, stromal-immune crosstalk, lineage trafficking, and feedback between cell states. Isolated cell culture and organoids provide mechanistic control but lose intact immune-organ architecture. *In vivo* animal experiments preserve systemic physiology but confound drug action with whole-body clearance, endocrine feedback, inter-organ crosstalk, and sparse sampling. A useful translational assay would sit between these extremes: controlled exposure in an intact immune organ, repeated sampling over pharmacodynamic time, and compatibility with cellular, transcriptomic, proteomic, and functional readouts.

The spleen is a logical organ for such a platform. It is the largest secondary lymphoid organ and a central site for blood filtration, innate sensing, antigen presentation, lymphocyte trafficking, and humoral organization.^4–7^ Its red pulp, white pulp, and marginal-zone architecture create immune functions that cannot be fully reproduced in reductionist culture systems. Earlier isolated spleen perfusion studies demonstrated that splenic viability, filtration, lymphocyte egress, and human spleen biology can be studied *ex vivo.*^8–14^ This platform is an attempt to modernize that concept into a contemporary multi-omic drug-testing platform.

Normothermic machine perfusion provides the engineering foundation. Originally developed for organ preservation and transplantation, *ex vivo* organ perfusion is increasingly being used for disease modeling, drug delivery, pharmacokinetics, biomarker discovery, and extended organ interrogation.^15–21^ In this setting, the effluent can function as a repeated organ-level liquid biopsy. Each sample contains soluble factors and cells exiting the organ, enabling kinetic measurements that are difficult to obtain *in vivo* and impossible to obtain from static endpoint tissue analysis.

A critical future extension is the human donor spleen. Spleens recovered during deceased-donor abdominal procurement are rarely transplanted and are most often discarded. With appropriate consent, organ-procurement-organization partnerships, and rapid recovery logistics, these tissues could become a high-value source of biologic information rather than waste material. A standardized human-spleen perfusion assay could generate donor-resolved immune-pharmacology datasets across age, sex, comorbidity, infection history, medication exposure, cause of death, and transplant-relevant immune states.

Here, we establish acellular normothermic porcine spleen perfusion as a translational immune-organ assay and use methylprednisolone as an intentionally stringent pharmacologic stress test. Glucocorticoids were selected because their acute immunosuppressive biology is clinically familiar, rapidly inducible, and measurable through both gene-regulatory and protein-level mechanisms. We therefore asked whether serial effluent sampling could recover expected steroid-responsive transcriptional programs, validate a nominated transcriptional marker (SOCS3) at the protein level, and expose non-transcriptional protein dynamics that would motivate follow-up human-cell assays.

## Results

### An acellular normothermic spleen perfusion platform for serial immune pharmacodynamics

We positioned acellular normothermic spleen perfusion as an organ-level translational checkpoint between reductionist preclinical testing and patient studies (Fig. 1A). The circuit is a recirculating extracorporeal loop in which an intact porcine spleen is cannulated via the splenic artery and perfused with an albumin-based, hemoglobin-free crystalloid/DMEM perfusate at 36–37°C, supported by a roller pump, membrane oxygenator, heat exchanger, venous reservoir, and a hemofiltration/dialysis module that allows dynamic perfusate composition control. The acellular formulation was deliberate: it removes exogenous leukocytes from the readout, so that cells and cell-associated proteins recovered in the venous effluent originate from the perfused spleen itself. Drug is administered as a bolus into the arterial limb of the perfusion circuit, and serial venous-effluent sampling at Pre-A, Pre-B, 1, 3, 6, and 12 h post-bolus (Fig. 1B) yields cellular material for paired RNA-sequencing and DIA mass-spectrometry proteomics from the same effluent stream. The design is analytical rather than systemic: the circuit exposes the intact splenic vascular bed to a defined perturbation and converts the organ response into a longitudinal molecular and cellular pharmacodynamic profile.

**Fig. 1.**
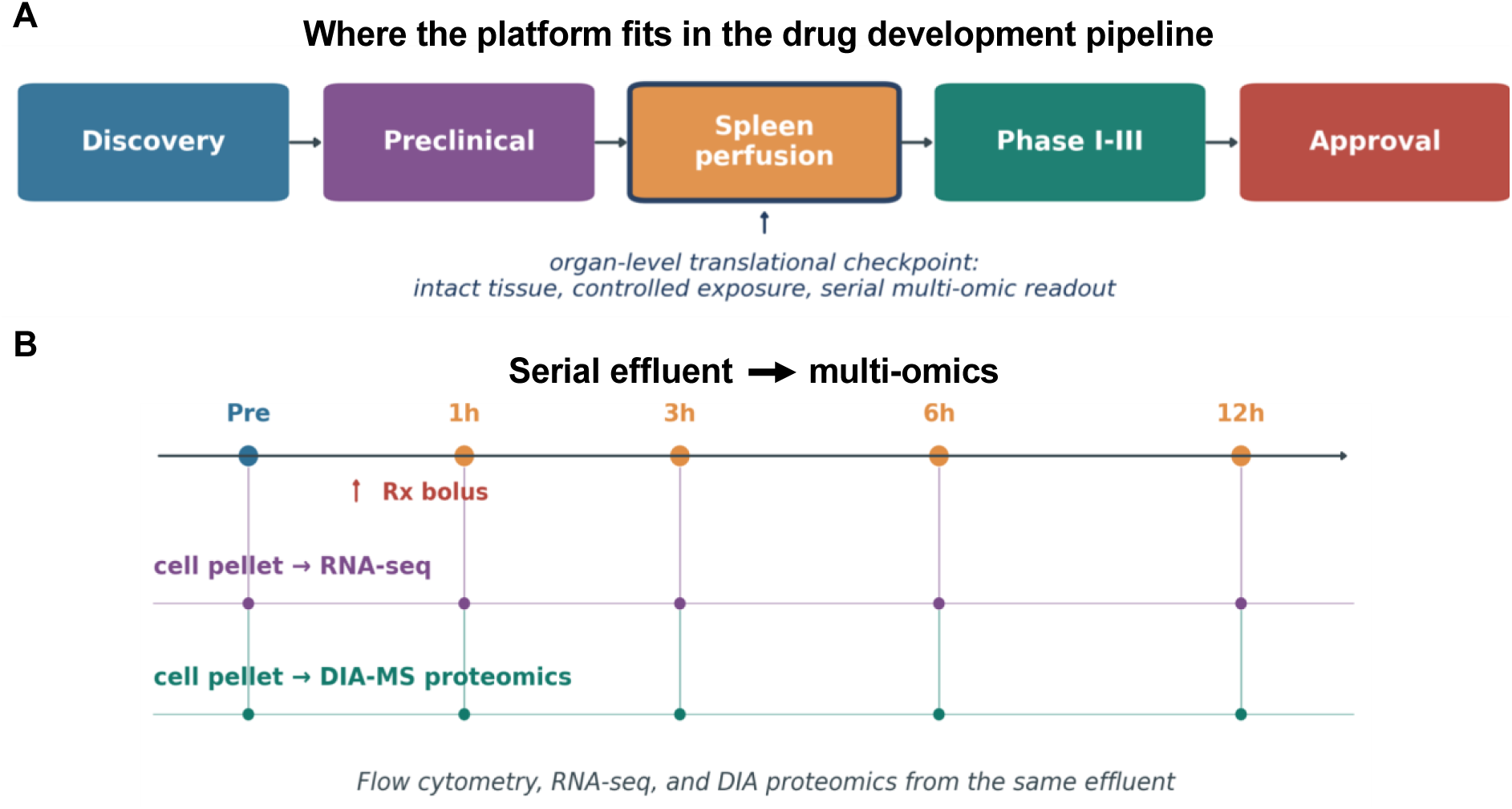
Acellular normothermic spleen perfusion as an analytical drug-testing platform. (**A**) The platform sits between conventional preclinical testing and Phase I-III clinical trials as an organ-level translational checkpoint, providing intact tissue, controlled drug exposure, and serial multi-omic readout. (**B**) Serial-effluent sampling design: pre-bolus baseline plus four post-bolus timepoints (1, 3, 6, 12 h) feeding both bulk RNA-sequencing and DIA mass-spectrometry proteomics from the same effluent stream, enabling kinetic integration of cellular, transcriptomic, and proteomic responses to the perturbation.

### Acellular spleen perfusion supports repeated viable-cell recovery and longitudinal flow analysis

Across four independent porcine perfusions on this circuit, the spleen maintained an aerobic metabolic envelope across the 12-hour window (Fig. 2A). Oxygen consumption (VO_2_) tracked between approximately 1.7 and 3.0 mL/min, oxygen delivery (DO_2_) ranged from approximately 4 to 8 mL/min, and perfusate pH remained between 7.18 and 7.55 throughout the run. Lactate accumulated from a baseline near 0.3 mmol/L to approximately 9.7 mmol/L at 12 h — a measurable but interpretable buildup characteristic of acellular metabolic operation, in which lactate clearance by extra-splenic tissues is absent by design. Histological inspection of the cannulated spleen at 1, 3, 6, and 12 h post-bolus confirmed preservation of red and white pulp microarchitecture, with intact lymphoid follicles visible at every post-treatment timepoint. (Fig. 2B-C)

**Fig. 2.**
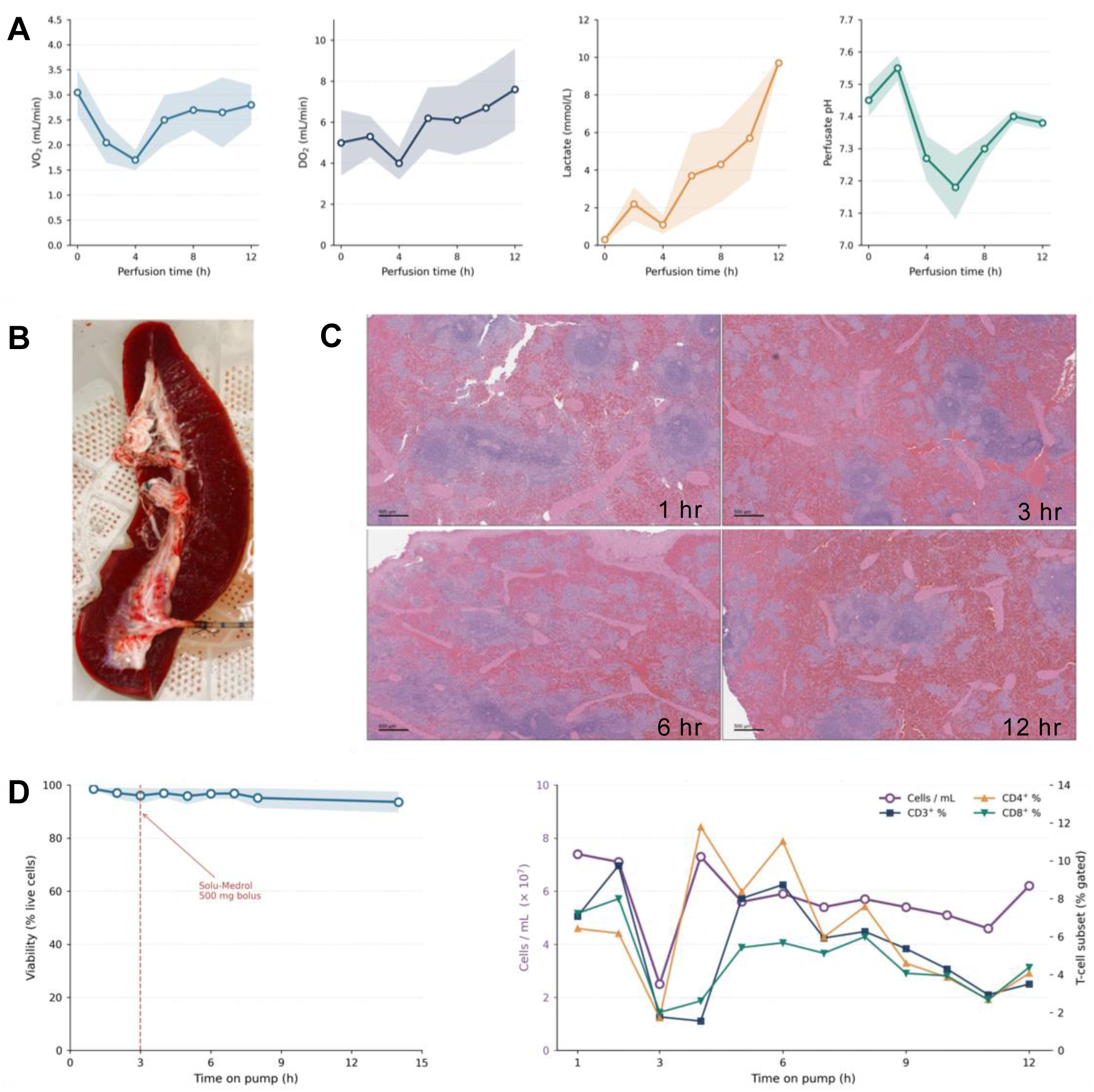
Empirical validation of the acellular spleen-perfusion platform. **A**) Hemodynamic and metabolic dynamics across n = 4 independent acellular perfusions (mean ± SEM, 2-h binning): oxygen consumption (VO_2_), oxygen delivery (DO_2_), perfusate lactate, and perfusate pH. (**B**) Representative photograph of an intact porcine spleen mounted on the perfusion circuit, showing splenic-artery cannulation and capsular preservation in the basin/sling containment system. (**C**) H&E-stained spleen biopsy sections at 1, 3, 6, and 12 h post-bolus (4× original magnification, scale bars = 500 µm), confirming preservation of red and white pulp microarchitecture across the perfusion window. (**D**) Flow cytometric monitoring of cellular output. Left: mean SYTOX-defined viability across two independent acellular perfusions over 14 h on pump; the shaded band shows the inter-perfusion range, and the vertical dashed line marks the 500 mg Solu-Medrol bolus at hour 3. Right: total effluent cellularity (cells/mL, plum) and CD3^+^ / CD4^+^ / CD8^+^ subset percentages across the 12-h effluent series of an independent acellular perfusion.

Flow cytometry across two independent acellular perfusions documented sustained viability throughout extended perfusion (Fig. 2D, left): SYTOX-defined live fractions remained between approximately 90% and 99% across 14 hours on pump, with the inter-perfusion range narrow throughout. In the paired CD3/CD4/SYTOX and CD3/CD8/SYTOX panels, live fractions ranged from 92.3% to 97.8% and 91.6% to 96.0%, respectively, and CD3^+^ cells remained measurable after steroid challenge — between 1.37% and 2.30% of live cells in the CD3/CD4 panel and 1.15% to 1.93% in the CD3/CD8 panel — with CD4/CD3 and CD8/CD3 fractions of 29%–64% and 35%–50%, respectively. Hourly effluent sampling from a separate acellular perfusion provided complementary cellularity and T-cell-subset readouts (Fig. 2D, right): total cellularity remained between approximately 2.5 and 7.4 × 10^7^ cells per mL, and CD3^+^, CD4^+^, and CD8^+^ subsets remained reliably detectable across the 12-hour series. These data do not on their own prove preserved organ function, but they show that the platform reliably recovers viable cells suitable for flow cytometry and downstream omics an analytical requirement for a serial immune-organ pharmacology assay.

### Serial effluent RNA-seq identifies steroid-responsive transcriptional programs and nominates SOCS3

To test whether the platform could detect acute immunosuppressive pharmacology, methylprednisolone sodium succinate was administered as a single bolus after baseline sampling. RNA-seq was performed on cells recovered from serial venous effluent. Targeted visualization of mean CPM values showed rapid induction of canonical glucocorticoid-response transcripts, including DUSP1, FKBP5, PER1, DDIT4, SGK1, KLF9, and ANXA1 (Fig. 3A).^22–27^ These early transcripts served as an internal positive-control module: cells exiting the perfused spleen reported steroid exposure through gene programs that are well established in glucocorticoid biology. The same dataset resolved pathway-level feedback programs that are central to steroid immunosuppression. NFKBIA and TNFAIP3 increased early, consistent with induction of negative regulators of NF-κB signaling; downstream inflammatory components showed later decline or mixed remodeling (Fig. 3B). JAK/STAT suppression was captured most clearly through SOCS-family induction. SOCS1, SOCS3, and CISH were induced early after methylprednisolone exposure, followed by broader changes across JAK, STAT, IRF, and interferon-associated transcripts (Fig. 3C). Because SOCS3 had one of the strongest early transcriptional responses and is a proximal suppressor of cytokine-JAK/STAT signaling, we selected it for orthogonal validation. In CD3/CD28-activated primary murine splenocytes, prednisone increased SOCS3 protein abundance by western blot at 3 h (Fig. 3D), converting the effluent RNA-seq observation into an independent protein-level pharmacodynamic validation. The data show that serial effluent RNA-seq can recover known glucocorticoid transcriptional programs and nominate validated pharmacodynamic markers from an intact immune organ. ^28–30^

**Fig. 3.**
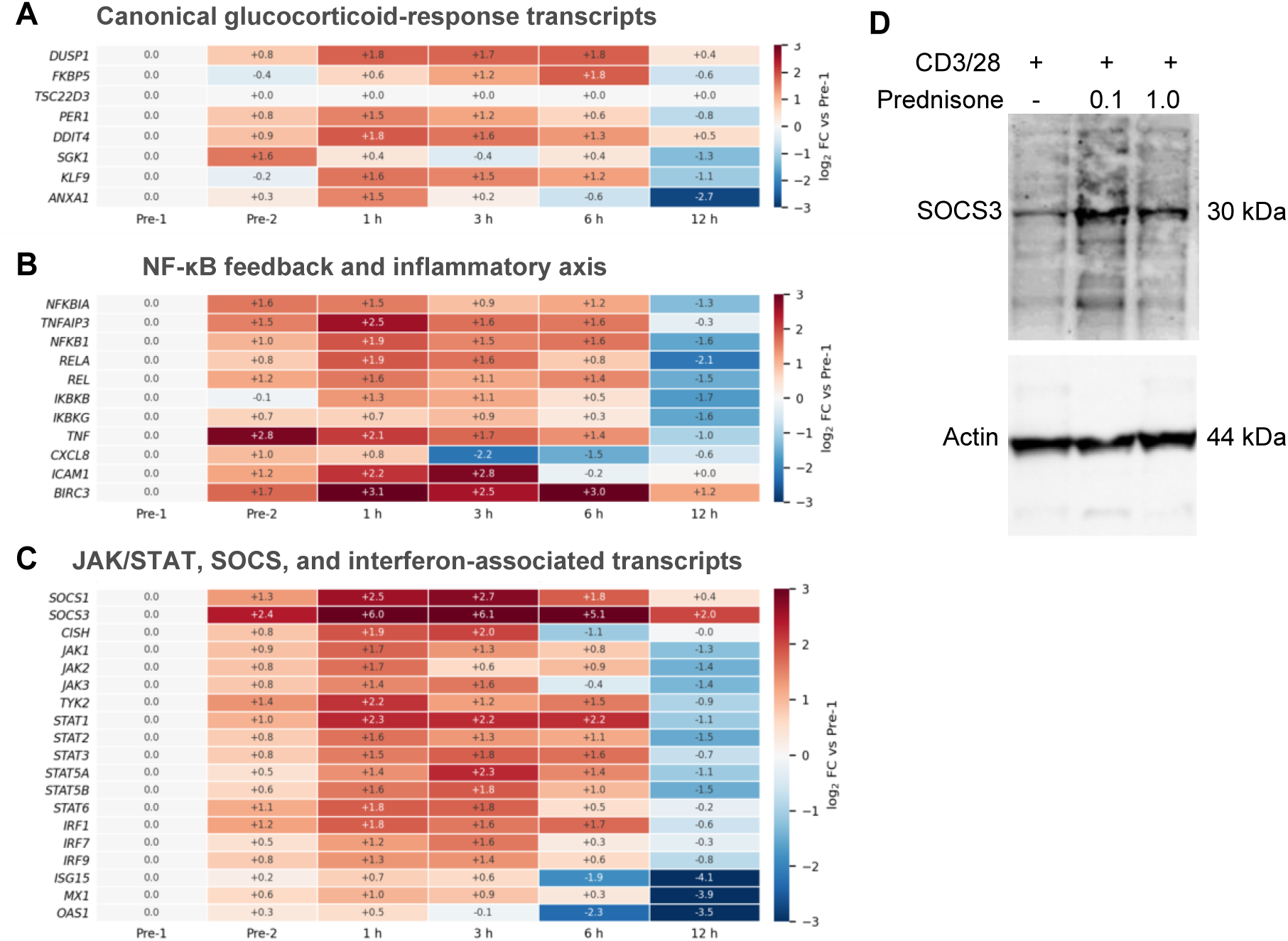
Serial effluent RNA-seq identifies glucocorticoid-responsive transcriptional programs and supports SOCS3 protein validation. (A) Canonical glucocorticoid-response transcripts show rapid induction after methylprednisolone exposure. (B) NF-κB feedback and inflammatory transcripts show early feedback-regulator induction and later pathway remodeling. (C) JAK/STAT, SOCS, and interferon-associated transcripts show early SOCS-family induction followed by broader pathway changes. Values are descriptive mean CPM log2 fold-changes relative to the Pre-1 baseline; color scale capped at ±3 for display. n = 3 technical replicates per timepoint from a single perfusion. (D) Primary murine splenocytes were stimulated in vitro with anti-CD3/CD28 and treated with prednisone (0.1 or 1.0 µM) for 3 h before harvest. Western blotting demonstrated increased SOCS3 protein abundance after prednisone treatment, providing an orthogonal validation of the SOCS3 signal nominated by serial effluent RNA-seq.

### Effluent-cell proteomics nominates a non-transcriptional CLC/Galectin-10-like response

DIA proteomics was performed on pelleted effluent cells from one representative perfusion in technical triplicate to determine whether the same serial effluent stream could identify protein-level pharmacodynamic signals that were not reducible to contemporaneous transcript abundance. Because the analyzed material was a cell pellet rather than a separately clarified soluble supernatant, these data are presented as effluent-cell-associated protein abundance rather than definitive soluble secretion.

The proteomic readout identified dynamic immune-associated protein groups after methylprednisolone exposure (Fig. 4). The most prominent galectin-family signal was a Sus scrofa protein group carrying an LGALS13 database annotation (Fig. 4A). We treat this as a porcine orthologous CLC/Galectin-10-like galectin rather than as canonical human placental Galectin-13/PP13 biology. This distinction is important: the database symbol provides the porcine locus label, whereas the translational assay question is whether the human immune counterpart of this protein axis has T-cell activity. The corresponding transcript was undetectable across the RNA-seq matrix, whereas other galectin transcripts such as LGALS1, LGALS3, LGALS8, and LGALSL were detectable. Thus, the signal nominated a non-transcriptional mechanism: cell-associated protein abundance changed without a matched effluent-cell transcript, suggesting either pre-existing protein mobilization, release from a specific cell subset, selective cell egress, or post-transcriptional regulation. The proteomic data also showed divergent immune-lineage-associated kinetics (Fig. 4B). MS4A1/CD20 remained elevated across the sampled post-steroid intervals, whereas several T-cell-associated and cytotoxic markers (CD3E, CD4, LCK, GZMB, GNLY) declined by 12 h or became undetected in the exploratory dataset; PTPRC/CD45 declined modestly. Selected NF-κB and JAK/STAT-axis protein groups showed dynamics broadly consistent with the transcriptional program, including early JAK3 elevation and late STAT3 decline (Fig. 4C).

**Fig. 4.**
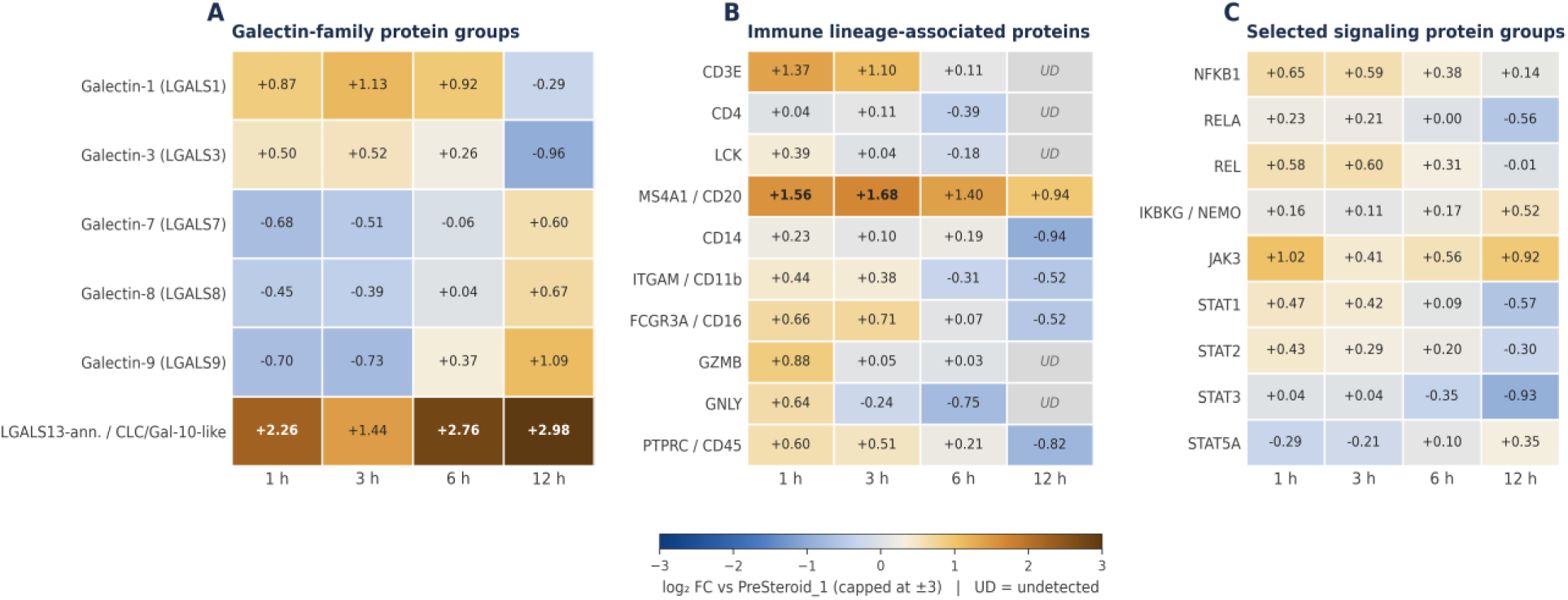
Exploratory effluent-cell proteomics nominates non-transcriptional protein-level steroid-response dynamics. (A) Galectin-family protein groups, including an LGALS13-annotated / CLC-Galectin-10-like porcine orthologous protein group. (B) Immune lineage-associated proteins (T-cell, B-cell, myeloid, cytotoxic, and pan-leukocyte markers). (C) Selected signaling protein groups (NF-κB and JAK/STAT axes). DIA proteomics was performed on effluent cell pellets from one representative biological perfusion in technical triplicate and should be interpreted as exploratory cell-associated protein abundance. Values are log2 fold-changes relative to the Pre-1 baseline; heatmap scale capped at ±3 for display. UD = undetected.

### Human Galectin-10 drives apoptosis in Jurkat T cells

The human Galectin-10 Jurkat assay was performed because the porcine DIA hit was annotated as LGALS13 but interpreted as an orthologous CLC/Galectin-10-like signal with a plausible human immune counterpart. We therefore used recombinant human Galectin-10 to test whether the protein could directly affect activated T cells, while including recombinant human Galectin-13 as a related galectin comparator. This assay was designed as an orthogonal functional follow-up module, not as proof that the porcine protein identity is fully resolved or that galectins causally drive the perfusion phenotype. CD3/CD28-stimulated Jurkat cells were treated with recombinant human Galectin-10 or Galectin-13 at 20 µg/mL for 24 h, and apoptosis/cell death was measured by Annexin V and propidium iodide (PI) staining.^31–32^ Recombinant human Galectin-10 produced a marked increase in Annexin V positivity and Annexin V/PI double-positivity. Annexin V+ cells increased from 8.0 ± 0.3% in resting cells and 19.4 ± 0.9% in stimulated controls to 84.3 ± 7.2% with human Galectin-10. Annexin V+ / PI+ double-positive cells increased from 3.7 ± 0.3% in resting cells and 12.1 ± 0.5% in stimulated controls to 80.7 ± 7.0% with human Galectin-10 (Fig. 5). Galectin-13 produced a more modest increase.

**Fig. 5.**
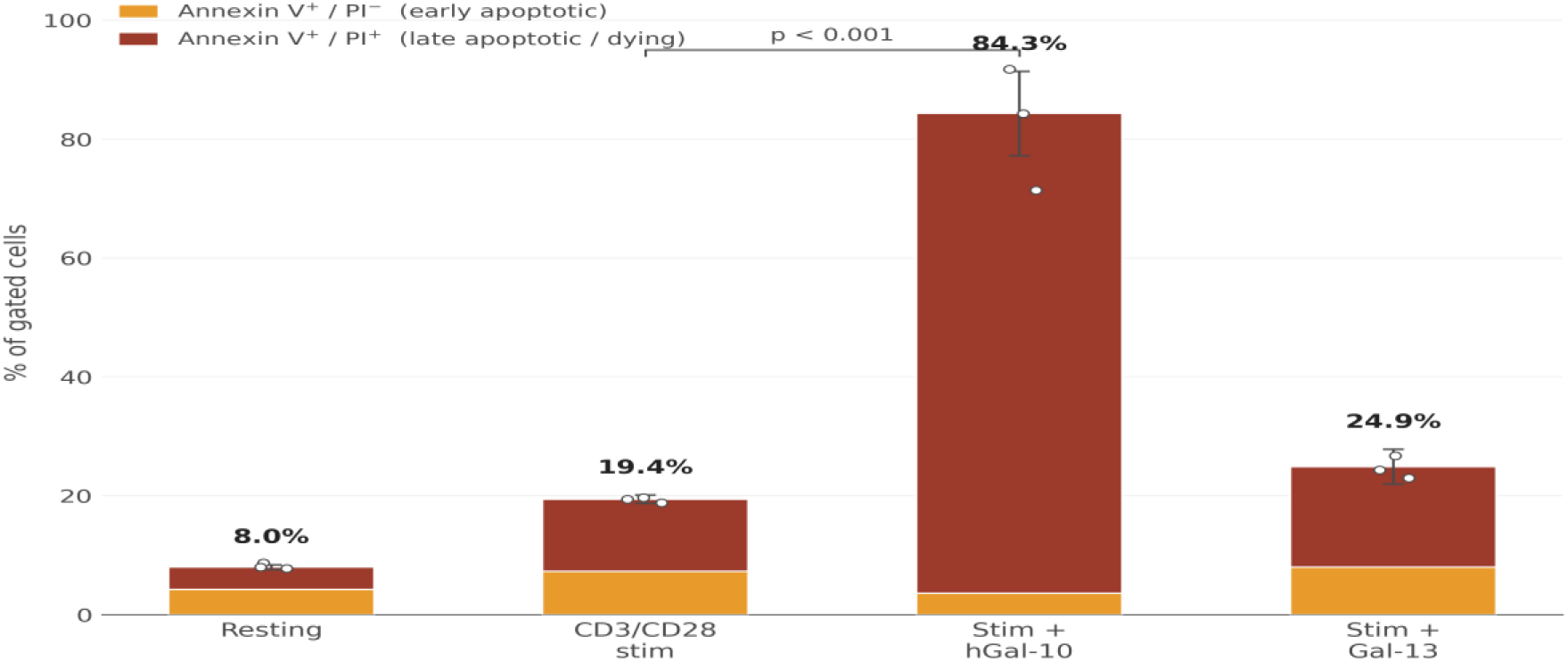
Human recombinant Galectin-10 induces high apoptosis in CD3/CD28-stimulated Jurkat T cells. Jurkat T cells were stimulated with anti-CD3/CD28 (5 µg/mL) for 1 h before addition of recombinant human Galectin-10 or Galectin-13 (20 µg/mL) for 24 h. Cells were stained with Annexin V and propidium iodide (PI) and analyzed by flow cytometry. Stacked bars decompose the total Annexin V+ population into early-apoptotic (Annexin V+ / PI−, gold) and late-apoptotic / dying (Annexin V+ / PI+, red) fractions. Total Annexin V+ values are annotated above each bar; error bars represent SEM on the total. Total Annexin V+ values were compared across conditions by one-way ANOVA with Dunnett’s multiple-comparison test against the stimulated control. Individual replicate values are shown as overlaid points (n = 3 per condition). Human Galectin-10 produced ∼84% total Annexin V+ positivity, the great majority of which had progressed to the late-apoptotic / dying state by 24 h compared with stimulated alone (p<0.001). Galectin-13 produced a more modest increase. This assay was selected because the porcine LGALS13-annotated proteomic signal is interpreted as a CLC/Galectin-10-like orthologous axis; it supports prioritization of that axis but does not prove porcine protein identity or causality in the perfused spleen.

### Cross-species comparison supports human donor-spleen deployment

To place the porcine signature in a translational frame, we cross-referenced direction-matched glucocorticoid response indices against published methylprednisolone responses in primary human leukocytes (GSE112101). At three classical glucocorticoid targets (NFKBIA, DUSP1, FKBP5), the porcine perfusion signature was concordantly induced with all four human leukocyte populations (Fig. 6A). SOCS1 was concordantly induced in B cells, CD4+ T cells, and monocytes and in the porcine spleen, with mild discordance in neutrophils. By contrast, SOCS3 and CISH were robustly induced in the porcine whole-organ signature (∼+4.5 and ∼+1.5 log2, respectively, at 1–3 h) but were either flat or repressed in three of four human leukocyte populations. SOCS3 is therefore more than a positive-control transcript in this study: it is a validated pharmacodynamic marker that exposes a whole-organ versus peripheral-blood difference. That discordance may reflect tissue-resident cell populations, lineage trafficking, or organ-specific feedback not represented in purified peripheral-blood leukocyte datasets.

**Fig. 6.**
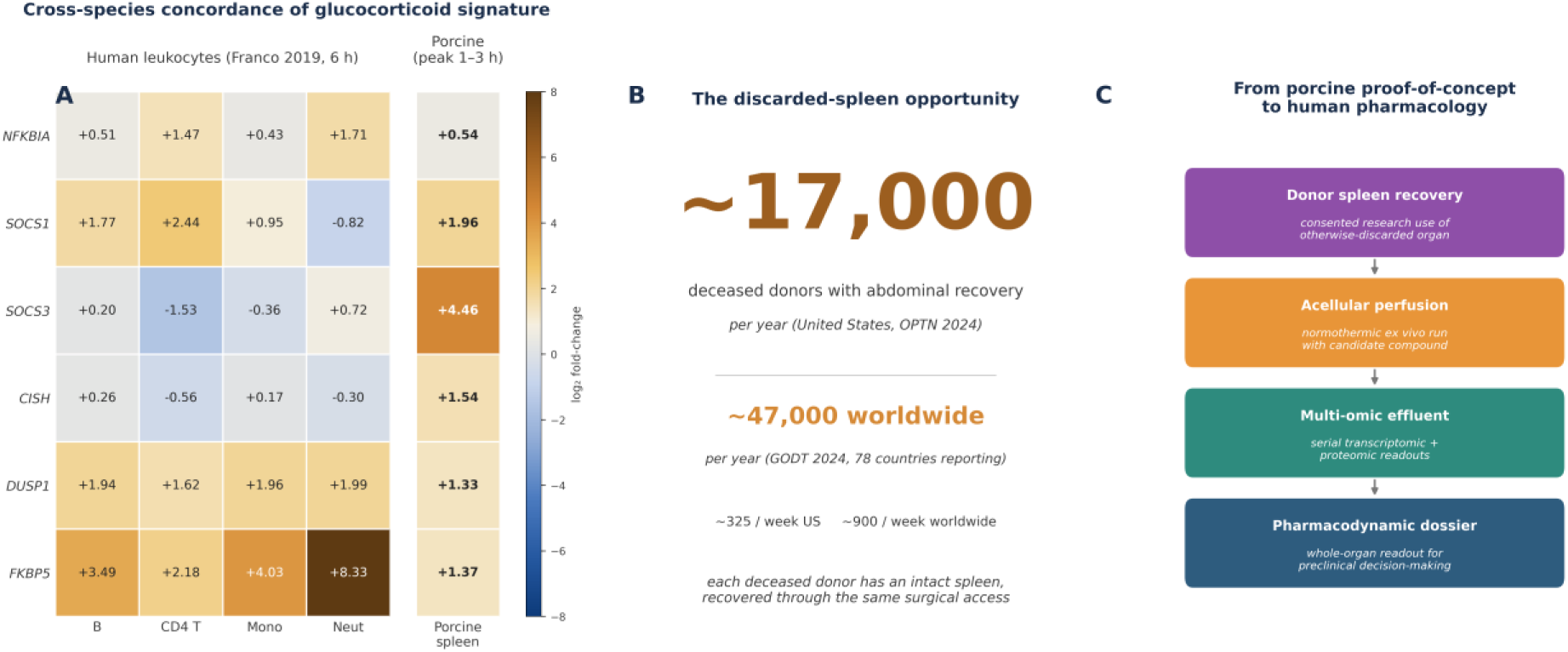
Cross-species comparison and the discarded-donor-spleen translational opportunity. (A) Heatmap of log2 fold-changes for six glucocorticoid-responsive genes across four primary human leukocyte populations (B cells, CD4+ T cells, monocytes, neutrophils) at 6 h methylprednisolone treatment from Franco et al. (23, GSE112101), alongside the present porcine spleen perfusion (peak induction at 1–3 h post-bolus). NFKBIA, DUSP1, and FKBP5 are concordantly induced across both species and all human cell types. SOCS1 is concordantly induced in three of four human populations and in the porcine spleen. SOCS3 and CISH are robustly induced in the porcine whole-organ signature but show cell-type-specific or repressive responses in purified human leukocyte populations; SOCS3 was further supported by independent protein-level validation in activated splenocytes. (B) Approximately 17,000 deceased donors with abdominal organ recovery occur each year in the United States (OPTN 2024; ∼325 per week), and ∼47,000 worldwide (GODT 2024, 78 countries reporting; ∼900 per week). Because the spleen is recovery-accessible through the same surgical exposure used for routine kidney, liver, and pancreas procurement, it could be co-recovered under appropriate research consent at no additional surgical cost — an untapped immunopharmacology resource that the platform is designed to convert into multi-omic pharmacodynamic data. (C) Proposed translational workflow: donor spleen recovery → acellular perfusion with candidate compound → multi-omic effluent profiling → pharmacodynamic dossier for preclinical decision-making.

The broader output of the study is an analytical architecture rather than a single biomarker. Serial spleen perfusion can generate physiologic, cellular, transcriptomic, proteomic, and functional data from the same organ-level perturbation. That creates a decision product for translational development: candidate ranking, organ-level activity, lineage-associated toxicity, pharmacodynamic-marker nomination, non-transcriptional mechanism discovery, and rational selection of combinations or dosing windows. The most powerful next iteration is human (Fig. 6B, C). Spleens from deceased donors are often discarded because they are not routinely transplanted. Under appropriate consent and recovery workflows, these organs could support short-duration human ex vivo perfusion studies that are impossible to perform in living patients and more clinically grounded than animal models alone. A donor-resolved human-spleen platform could test immunosuppressants, biologics, cell-targeting agents, nanoparticles, checkpoint modulators, inflammatory-pathway inhibitors, and AI-prioritized candidates against intact human immune-organ architecture.

## Discussion

This study establishes acellular normothermic spleen perfusion as a serial immune-organ assay that can connect controlled pharmacologic exposure to transcriptional programs, protein-level mechanisms, and orthogonal functional validation. The central translational claim is not that a porcine perfusion reproduces human immunity, but that intact-organ perfusion can generate an interpretable pharmacodynamic data prior patient exposure.

Serial sampling recovered consistently viable cells throughout the run. Across multiple independent perfusions and viability was maintained throughout the 14-hour operating window. The methylprednisolone experiment then served as a pharmacologic positive control and a stress test of mechanistic resolution. RNA-seq captured rapid induction of expected glucocorticoid-responsive genes and feedback inhibitors, including DUSP1, FKBP5, NFKBIA, SOCS1, SOCS3, and CISH. SOCS3 deserves particular emphasis because it was not left as an effluent transcript alone: the protein-level increase in prednisone-treated, CD3/CD28-activated primary murine splenocytes supports SOCS3 as a candidate pharmacodynamic marker for glucocorticoid action. The ability to resolve the timing of these responses is a core advantage of the platform; it provides a kinetic pharmacodynamic signature rather than a static endpoint measurement.

DIA proteomics provided a second class of signal: a non-transcriptional protein-level candidate mechanism of steroid immunosuppression. The most prominent galectin-family protein group carried a Sus scrofa LGALS13 annotation but is treated here as a porcine orthologous signal to human CLC/Galectin-10 for translational testing. That annotation drove the in vitro design: recombinant human Galectin-10 was selected because it is the human immune galectin corresponding to this CLC-like axis, and Galectin-13 was included as a related comparator. In activated Jurkat cells, human Galectin-10 produced marked Annexin V and Annexin V/PI positivity at 24 h.

## Limitations

There are several limitations of the manuscript. First, the current perfusate is acellular and lacks erythrocytes, complement, plasma proteins, circulating leukocyte input, endocrine feedback, and systemic clearance; this makes the assay analytically clean but biologically incomplete. Second, the key proteomics dataset comes from one biological perfusion with technical triplicates and should be considered exploratory until validated across independent organs. Third, late timepoints occur in the setting of rising lactate which may interfere with the transcriptional response. Fourth, the SOCS3 validation was performed in activated murine splenocytes rather than in post-perfusion porcine effluent cells, so it validates steroid-induced SOCS3 protein induction but not cell-type-specific causality in the perfused spleen. Fifth, the Jurkat Galectin-10 assay is a cell-line functional screen, not a primary-cell validation and not a causal test of the perfused-spleen response. Finally, species-specific considerations apply to galectin-family annotations in the porcine reference proteome, and the LGALS13/CLC-Galectin-10-like orthology assignment will require peptide-level confirmation for translation to human spleen. Eighth, the 12-hour operating window captures acute pharmacodynamics but does not address tolerance induction or long-term immune reconstitution.

## Outlook

The next potential experiments are straightforward and high value: Human donor spleens that would otherwise be discarded could become a renewable source of clinically anchored immune-organ pharmacology. In the AI drug-development era candidate generation is accelerating, but translation still fails when biology is measured in systems that are too simple or too late. Ex vivo human spleen perfusion could provide an intermediate checkpoint — not a replacement for animals or clinical trials, but a high-information bridge that filters, ranks, and mechanistically annotates immune-active candidates before they reach patients.

## Materials and Methods

### Ethical approval and animal tissue use

Porcine tissue from White Yorkshire x Landrace pigs (weight 40-50 kg, 2 males, 2 females) ordered from Oakhill Genetics (Ewing, IL) was obtained under Institutional Animal Care and Use Committee (IACUC) approval at the Arkansas Children’s Research Institute (Animal Welfare assurance number: A3063-01). Spleens were procured from animals euthanized for purposes unrelated to this study. No animals were sacrificed solely for this research.

### Organ procurement and acellular normothermic perfusion

Porcine spleens were procured using standard surgical techniques, flushed via the splenic artery with cold isotonic solution, and maintained cold until cannulation. The splenic artery was cannulated and the organ was placed in a low-compression hybrid containment system consisting of a rigid basin and compliant sling support, designed to minimize capsular compression, allow unrestricted venous outflow, and permit continuous visual monitoring. Perfusion was performed through a recirculating extracorporeal circuit comprising a roller pump, membrane oxygenator, heat exchanger, venous reservoir, and hemofiltration/dialysis module. The acellular perfusate included balanced crystalloid, albumin (5–25%), electrolytes, sodium bicarbonate, glucose, and DMEM. No hemoglobin-based oxygen carrier was used. Perfusion was conducted at 36–37 °C under flow-controlled conditions with passive venous drainage.

### Drug intervention and serial sampling

After stabilization and baseline sampling (Pre-A and Pre-B), methylprednisolone sodium succinate (Solu-Medrol; 500 mg for the multi-omic run unless otherwise specified) was administered as a single circuit bolus into the arterial limb of the perfusion circuit. Serial venous-effluent samples were collected at 1, 3, 6, and 12 h post-bolus. Each multi-omic timepoint contributed three independent technical replicates (A, B, C) drawn from the recirculating effluent within minutes of one another.

### Acellular perfusion flow cytometry and physiologic monitoring

For the two acellular perfusions used to characterize cellular viability, serial samples were analyzed using SYTOX viability dye and CD3/CD4/SYTOX or CD3/CD8/SYTOX panels with an intended acquisition target of approximately 500,000 events per panel. Steroid exposure (Solu-Medrol, 500 mg) was administered at hour 3 on pump in one of the two perfusions; the second perfusion provided an additional SYTOX viability series through 14 h on pump. These flow data were used as platform validation and were not treated as formal biological replicates. A separate independent acellular perfusion provided hourly effluent cellularity (cells/mL) and CD3^+^ / CD4^+^ / CD8^+^ subset percentages across the 12-hour series. Hemodynamic and metabolic parameters (VO_2_, DO_2_, O_2_ER, lactate, pH, PaO_2_, PvO_2_) were measured continuously across four independent perfusions and binned into 2-hour intervals (mean ± SEM). Splenic biopsies for histological evaluation were fixed, paraffin-embedded, stained with hematoxylin and eosin (H&E), and imaged at 4× magnification.

### RNA sequencing and descriptive transcriptomic analysis

Cells from serial effluent samples were pelleted, washed, and resuspended in Zymo DNA/RNA Shield, then shipped at room temperature to Plasmidsaurus for RNA extraction, library preparation, and Illumina sequencing. Reads were aligned to the *Sus scrofa* reference genome, and transcript abundance was summarized as Counts Per Million (CPM). For the descriptive heatmaps shown here, timepoint-level mean CPM values were converted to log_2_ fold-change relative to the Pre-1 baseline using a uniform pseudocount. Future formal differential-expression analysis should use raw counts with a replicate-aware model such as DESeq2, edgeR, or limma-voom.

### DIA proteomics (exploratory effluent-cell-associated)

For DIA proteomics, 1 mL of cellular perfusate was pelleted, washed in PBS, frozen, and submitted to the UAMS Proteomics Core.

#### Orbitrap Astral DIA – 40 minutes - CME

Total protein from each sample was reduced, alkylated, and purified by chloroform/methanol extraction prior to digestion with sequencing grade modified porcine trypsin (Promega).

Tryptic peptides were then separated by a reverse phase Ion-Opticks-TS analytical column (25 cm x 75 um with 1.7 um C18 resin) supported by an EASY-Spray nano-source and stabilized with a Heater THOR Controller (Ion-Opticks) at 60°C. Peptides were trapped and eluted from a (PepMap Neo, 300um x 5mm Trap) using a Vanquish Neo UHPLC nano system (Thermo Scientific) which kept the samples at 11°C before injection. Peptides were eluted at a flow rate of 0.350uL/min using a 35 min gradient from 98% Buffer A:2% Buffer B to 94.5:5.5 at 0.1 minutes to 56:44 at 27.1 minutes followed by a column wash of 45:55 at 29.7 minutes to 1:99 at 35 minutes followed by equilibration back to 98:2. Eluted peptides were ionized by electrospray (2.5 kV) followed by mass spectrometric analysis on an Orbitrap Astral mass spectrometer (Thermo). Precursor spectra were acquired from 380-980 Th, 240,000 resolution, normalized AGC target 200%, maximum injection time 3 ms. DIA acquisition on the Orbitrap Astral was configured to acquire 199, 3 Th window from 380-980 Th, 25% HCD Collision Energy, normalized AGC target 100%, maximum injection time 3 ms. Fragment (MS2) scan range from 150-2000 Th with an RF Lens (%) set to 40.

Buffer A = 0.1% formic acid, 0.5% acetonitrile in water

Buffer B = 80% acetonitrile, 20% water, 0.1% formic acid

#### Data Analysis: Spectronaut – VSN

Following data acquisition, data were searched using Spectronaut (Biognosys version 20.5) against the UniProt *Sus scrofa* database (Proteome ID: UP000008227, Taxon ID: 8923, 1st version of 2026) using the directDIA method with an identification precursor and protein q-value cutoff of 1%, generate decoys set to true, the protein inference workflow set to maxLFQ, inference algorithm set to IDPicker, quantity level set to MS2, cross-run normalization set to false, and the protein grouping quantification set to median peptide and precursor quantity. Fixed Modifications were set to Carbamidomethyl (C) and variable modifications were set to Acetyl (Protein N-term), Oxidation (M). Protein MS2 intensity values were assessed for quality using ProteiNorm (Graw *et al.*). The data was normalized using VSN (Huber et al) and analyzed using proteoDA to perform statistical analysis using Linear Models for Microarray Data (limma) with empirical Bayes (eBayes) smoothing to the standard errors (Thurman et al, Ritchie et al). Proteins with an FDR adjusted p-value < 0.05 and a fold change > 2 were considered significant.

Proteomics was performed on one representative biological perfusion in technical triplicate and is presented as exploratory effluent-cell-associated protein abundance rather than formal biological inference; the analyzed material was a cell pellet, not a separately clarified soluble supernatant. The galectin-family signal annotated as LGALS13 in the Sus scrofa reference is reported as a Sus scrofa LGALS13-annotated, CLC/Galectin-10-like protein group. This orthologous interpretation was the rationale for testing recombinant human Galectin-10 in a functional assay, but peptide-level confirmation will be required before assigning definitive porcine-to-human identity.

### Recombinant human galectin Jurkat functional assay

Jurkat E6-1 T cells were maintained in RPMI-1640 supplemented with 10% FBS and 1% penicillin/streptomycin. The assay was designed as an orthogonal functional test of the human immune counterpart of the porcine LGALS13-annotated / CLC-Galectin-10-like proteomic signal. Cells were stimulated with anti-CD3/anti-CD28 stimulation cocktail for 1 h and treated for 24 h with recombinant human Galectin-10 or Galectin-13 (20 µg/mL each). Apoptosis and cell death were measured by Annexin V / propidium iodide (PI) staining and flow cytometry. Annexin V+ (early + late apoptotic) and Annexin V+ / PI+ (late-apoptotic / dying) populations were quantified relative to resting and stimulation-only controls. Data are shown as mean ± SEM from n = 3 samples per condition. This assay was used as an orthogonal functional prioritization experiment and was not interpreted as proof of porcine protein identity or causality in the perfused spleen.

### Cross-species comparison

Direction-matched human glucocorticoid log_2_ fold-changes for NFKBIA, SOCS1, SOCS3, CISH, DUSP1, and FKBP5 were extracted from the Franco *et al.*^29^ GSE112101 differential-expression results at the 6-h methylprednisolone timepoint for B cells, CD4^+^ T cells, monocytes, and neutrophils. Porcine perfusion values used peak induction (max of 1 h and 3 h post-bolus) for each gene. Both human and porcine values are plotted as descriptive log_2_ fold-changes; no formal cross-species statistical equivalence test was performed.

### Visualization and statistics

Figures were generated using Python and Matplotlib with a harmonized visual style across panels. Flow and Jurkat data are shown as descriptive mean ± SEM or raw sample points where appropriate. RNA-seq heatmaps are descriptive log_2_ fold-change visualizations relative to the Pre-1 baseline. Proteomics heatmaps show normalized log_2_ fold-change values from a single biological perfusion with technical triplicates; these data should be interpreted as hypothesis-generating.

### Statistical analysis

All statistical analyses were performed in GraphPad Prism v10 and Python 3.11 (scipy.stats, statsmodels). Continuous data are reported as mean ± SEM unless otherwise indicated. For the Jurkat galectin apoptosis assay (Fig. 5; n = 3 per condition), total Annexin V+ values were compared across conditions (resting, stimulated, stimulated + Galectin-10, stimulated + Galectin-13) by one-way ANOVA with Dunnett’s multiple-comparison test against the stimulated control; p < 0.05 was considered significant. Flow-cytometric viability and T-cell subset percentages from the acellular perfusion runs are reported as descriptive statistics (mean, SEM, and inter-perfusion range) without formal hypothesis testing, because the perfusions were treated as independent platform-validation runs rather than biological replicates of a single experimental condition. Hemodynamic and metabolic parameters across the four independent perfusions (VO2, DO2, lactate, pH) are reported as mean ± SEM in 2-hour bins, also as descriptive validation statistics. RNA-seq fold-changes presented in Figs. 3 and 6 are descriptive log2(mean CPM + pseudocount) values relative to the Pre-1 baseline; no per-gene differential-expression test was applied to the figure data, and replicate-aware count-based modeling (DESeq2, edgeR, or limma-voom) is identified in the Limitations as the appropriate next analytical step.

## Data availability

RNA-sequencing and proteomics accession numbers will be inserted after deposition and release.

## Use of AI-assisted tools

Large language model (LLM)-based AI tools were used to assist with data-analysis workflows, figure generation, manuscript editing, and spell-checking. All AI-assisted outputs were critically reviewed and verified by the authors, who assume full responsibility for the accuracy, integrity, and interpretation of all data, analyses, and conclusions presented. AI tools were not used to generate scientific hypotheses, experimental plans, or interpretation of experimental results.

## Acknowledgements

IDeA National Resource for Quantitative Proteomics and NIH/NIGMS grant R24GM137786.

Funding, author contributions, competing interests, and full data and materials availability statements will be completed at submission.

